# Elucidating the Impact of Bacterial Lipases, Human Serum Albumin, and FASII Inhibition on the Utilization of Exogenous Fatty Acids by *Staphylococcus aureus*

**DOI:** 10.1101/2023.06.29.547085

**Authors:** Emily L. Pruitt, Rutan Zhang, Dylan H. Ross, Nathaniel K. Ashford, Xi Chen, Francis Alonzo, Matthew F. Bush, Brian J. Werth, Libin Xu

**Affiliations:** Department of Chemistry, University of Washington, Seattle, Washington, USA; Department of Medicinal Chemistry, University of Washington, Seattle, Washington, USA; Department of Pharmacy, University of Washington, Seattle, Washington, USA; Department of Microbiology and Immunology, Loyola University Chicago-Stritch School of Medicine, Maywood, Illinois, USA

**Keywords:** *Staphylococcus aureus*, exogenous fatty acid, bacterial lipase, human serum albumin, AFN-1252, lipidomics

## Abstract

*Staphylococcus aureus* only synthesizes straight-chain or branched-chain saturated fatty acids (SCFAs or BCFAs) via the type II fatty acid synthesis (FASII) pathway, but as a highly adaptive pathogen, *S. aureus* can also utilize host-derived exogenous fatty acids (eFAs), including SCFAs and unsaturated fatty acids (UFAs). *S. aureus* secretes three lipases, Geh, sal1, and SAUSA300_0641, which could perform the function of releasing fatty acids from host lipids. Once released, the FAs are phosphorylated by the fatty acid kinase, FakA, and incorporated into the bacterial lipids. In this study, we determined the substrate specificity of *S. aureus* secreted lipases, the effect of human serum albumin (HSA) on eFA incorporation, and the effect of FASII inhibitor, AFN-1252, on eFA incorporation using comprehensive lipidomics. When grown with major donors of fatty acids, cholesteryl esters (CEs) and triglycerides (TGs), Geh was found to be the primary lipase responsible for hydrolyzing CEs, but other lipases could compensate for the function of Geh in hydrolyzing TGs. Lipidomics showed that eFAs were incorporated into all major *S. aureus* lipid classes and that fatty acid-containing HSA can serve as a source of eFAs. Furthermore, *S. aureus* grown with UFAs displayed decreased membrane fluidity and increased production of reactive oxygen species (ROS). Exposure to AFN-1252 enhanced UFAs in the bacterial membrane, even without a source of eFAs, indicating a FASII pathway modification. Thus, the incorporation of eFAs alters the *S. aureus* lipidome, membrane fluidity, and ROS formation, which could affect host-pathogen interactions and susceptibility to membrane-targeting antimicrobials.

**IMPORTANCE:** Incorporation of host-derived exogenous fatty acids (eFAs), particularly unsaturated fatty acids (UFAs), by *Staphylococcus aureus* could affect the bacterial membrane fluidity and susceptibility to antimicrobials. In this work, we found that Geh is the primary lipase hydrolyzing cholesteryl esters and, to a less extent, triglycerides (TGs) and that human serum albumin (HSA) could serve as a buffer of eFAs, where low levels of HSA facilitate the utilization of eFAs, but high levels of HSA inhibit it. The fact that the FASII inhibitor, AFN-1252, leads to an increase in UFA content even in the absence of eFA suggests that membrane property modulation is part of its mechanism of action. Thus, Geh and/or the FASII system look to be promising targets to enhance *S. aureus* killing in a host environment by restricting eFA utilization or modulating membrane property, respectively.

## INTRODUCTION

Antibiotic-resistant bacteria pose a major threat to global health, killing more people than HIV/AIDS or malaria (1). Among them, *Staphylococcus aureus* has been deemed one of the most serious threats, infecting the skin, soft tissue, and blood. It causes nearly 120,000 bloodstream infections with 20,000 associated deaths per year in the United States alone (2). *S. aureus* adapts to the host environment by incorporating exogenous fatty acids (eFAs) into its cell membrane, thereby allowing the bacteria to reduce energy consumption from *de novo* fatty acid biosynthesis, bypass the innate immune response, and withstand drug activity (3–11). Elucidating the effects of body fluids on the metabolism of the bacteria is critical to understanding the host-pathogen interaction and evolution of antimicrobial resistance (3, 12–14).

*S. aureus* only synthesizes straight-chain or branched-chain saturated fatty acids (SCFAs or BCFAs) via the type II fatty acid synthesis pathway (FASII) but can also utilize host-derived SCFA and unsaturated fatty acids (UFAs) or free fatty acids (FFA) (4, 6, 8, 15, 16). In our recent study, lipidomics analysis of *S. aureus* grown in human serum showed that bacteria incorporate UFAs into the bacterial membrane lipids, and cholesteryl esters and triglycerides are the major donors of fatty acid substrates in serum (3). Human serum albumin, an abundant carrier protein in the bloodstream that binds to acidic and lipophilic compounds, has been shown to sequester FFAs to restrict their exploitation by the bacteria (17, 18), but we hypothesize that it may also serve as a reservoir of fatty acids.

To facilitate the incorporation of eFAs into its membrane, *S. aureus* secretes three lipases, *S. aureus* lipase 1 (sal1), glycerol ester hydrolase (Geh), and SAUSA300_0641 (0641 or sal3), to release FFA from lipids found in serum (Figure 1) (4, 5, 15, 19–22). Once FFAs are released by the lipases, they can be taken up by the bacteria, phosphorylated by the fatty acid kinase (FakA), and incorporated into the bacterial lipids, with or without further elongation via the FASII pathway (Figure 1) (3–7). When using triglycerides (TGs) as substrates, Geh can release both short-(4-carbon) and long-chain substrates (16 and 18-carbon), with a preference for the long-chain fatty acids linoleic acid (18:2) and oleic acid (18:1), whereas sal1 prefers short-chain fatty acid (4-carbon) substrates (4, 5, 19–21). 0641 was also found to prefer hydrolyzing short-chain fatty acids (4-carbon or fewer) from triglycerides (22). Several studies have revealed the importance of these lipases as multifaceted virulence factors in *S. aureus* infections; however, the substrate specificity of Geh, sal1, and 0641 on cholesterol esters and the impact of Geh, sal1, or 0641 knockouts on eFA utilization have not been examined previously.

**Figure 1.**
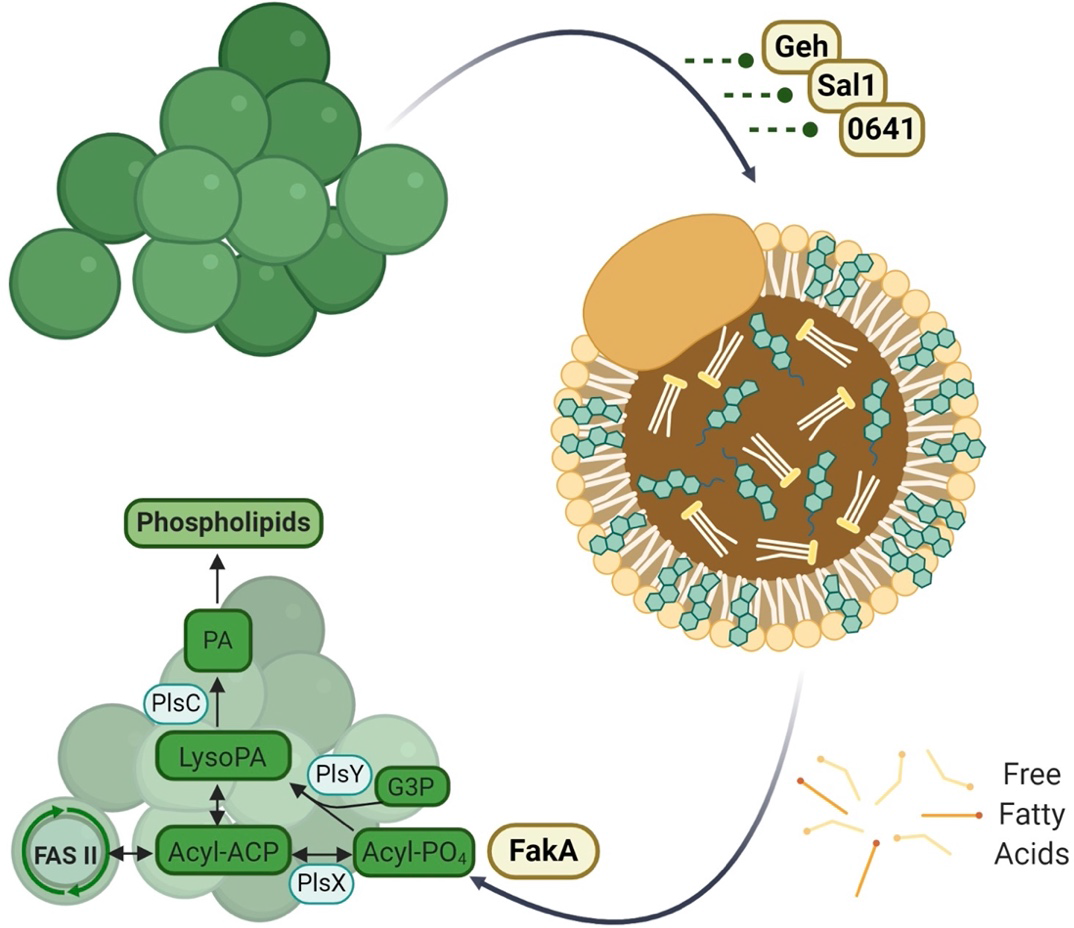
Schematic showing the release and utilization of exogenous fatty acids by *S. aureus*.

Incorporated serum UFAs can alter lipid packing, affecting the binding of membrane-targeting antimicrobials, and as an adaptive mechanism to drug exposure, *S. aureus* has been shown to modify its membrane and cell wall composition (9, 25–28). AFN-1252, a FabI inhibitor, has been developed as a FASII-targeting antibiotic, but its effect on broad lipidomic changes has not been well characterized (7). The therapeutic efficacy of AFN-1252 also remains in debate, as it shows promising treatment for skin and soft-tissue bacterial infections, but FASII bypassing variants that utilize host-derived eFAs, bring into question its long-term effectiveness (9–11, 18, 29–31). Although *S. aureus* can uptake eFAs and use them to evade the effects of FASII inhibitors and antibiotics, UFAs have also long been known to be toxic to the bacteria (4, 32–34). Polyunsaturated fatty acids (PUFAs), such as the abundant mammalian fatty acid arachidonic acid, can inflict damage on *S. aureus* upon incorporation into its membrane and kill the pathogen through a lipid peroxidation mechanism (32, 35, 36).

Despite previous work on the effect of exogenous fatty acids on *S. aureus,* several significant questions remain. First, the substrate specificity of the released lipases toward cholesterol esters remains unknown. Second, the comprehensive lipidomic changes resulting from eFA utilization have not been completely elucidated. Third, the role of albumin as a reservoir for fatty acids and its impact on eFA incorporation efficacy has not yet been determined. Fourth, the effect of the FASII inhibitor, AFN-1252, on eFA utilization has not yet been investigated. To answer these questions, we grew *S. aureus* and *geh*, *sal1*, *0641*, or *fakA* knockout (KO) mutant strains in tryptic soy broth (TSB) supplemented with eFAs under various conditions and conducted comprehensive lipidomic analyses of these bacteria. We further characterized the changes in membrane fluidity and formation of reactive oxygen species resulting from the incorporation of unsaturated eFAs.

We found that a) Geh is the primary lipase responsible for hydrolyzing cholesteryl esters, and, to a less extent, triglycerides; b) exogenous fatty acids were incorporated into the bacterial membrane when grown in serum regardless of the lipase knockout; c) human-serum albumin can serve as a buffer of eFA for *S. aureus*, facilitating the use of eFAs at a low concentration but inhibiting eFA utilization at high concentrations; d) AFN-1252 leads to an increase of UFAs in its membrane with or without eFAs; e) incorporation of unsaturated eFAs leads to increased membrane fluidity during the exponential growth phase; and f) incorporation of unsaturated eFAs increases reactive oxygen species formation, inhibiting *S. aureus* growth.

## RESULTS

### *S. aureus* lipase knockouts grown in serum still incorporate UFAs

*S. aureus* and *geh*-, *sal1*-, *0641*-, or *fakA*-knockout mutant strains (Δ*geh*, Δ*sal1*, Δ*0641*, or Δ*fakA*) were grown in the presence and absence of human serum, and changes in the lipidome were identified through hydrophilic-interaction liquid chromatography (HILIC) ion mobility-mass spectrometry (IM-MS) to determine the role of each enzyme in this environment. HILIC first resolves lipid species on a scale of seconds based on the polarity of the head groups and then by acyl chain length and degree of unsaturation within the subclass (37–39). Lipid separation is further increased through ion mobility, a gas-phase separation orthogonal to liquid chromatography (LC). As described previously, lipid identification is enhanced by using collisional cross section (CCS) values obtained from the IM-MS analysis (37, 38). Some serum-derived lipids, such as phosphatidylcholines, phosphatidylethanolamines, and sphingomyelins, are not truly incorporated into the bacterial membrane (3). Thus, major lipids that are synthesized by *S. aureus,* diglucosyldiacylglycerols (DGDGs), lysyl-phosphatidylglycerols (LysylPGs), phosphatidylglycerols (PGs), and cardiolipins (CLs), were examined (39–41).

We determined the total carbon and unsaturation degrees of the lipid acyl side chains for the major lipid species in the wild type (WT) and Δ*geh*, Δ*sal1*, Δ*0641*, Δ*fakA* mutants (Figure 2). As seen in the Figure, *S. aureus* grown in TSB-only displayed higher levels of fully saturated lipid species compared to those grown in human serum for each lipid class. This is not surprising since without exogenous fatty acids, the bacteria can only synthesize saturated SCFA or BCFA *de novo.* Consistent with previous studies, the Δ*fakA* mutant possessed a higher abundance of long acyl side chains (8). All strains contained DGDG, PG, and LysylPG saturated lipids with 32 to 37 total carbons, with 33 and 35 carbons being the major species across classes in each strain. Upon further targeted fragmentation experiments using tandem MS (MS/MS) on select DGDG and PG lipids, no differences in acyl chain composition were observed across the most abundant lipids of the wild type, Δ*geh*, Δ*sal1*, Δ*0641* and Δ*fakA* strains grown in TSB-only (Supplemental Data in Excel). C15:0 was consistently identified as the major component of saturated PGs while C20:0 was the most abundant FA moiety in DGDGs.

**Figure 2.**
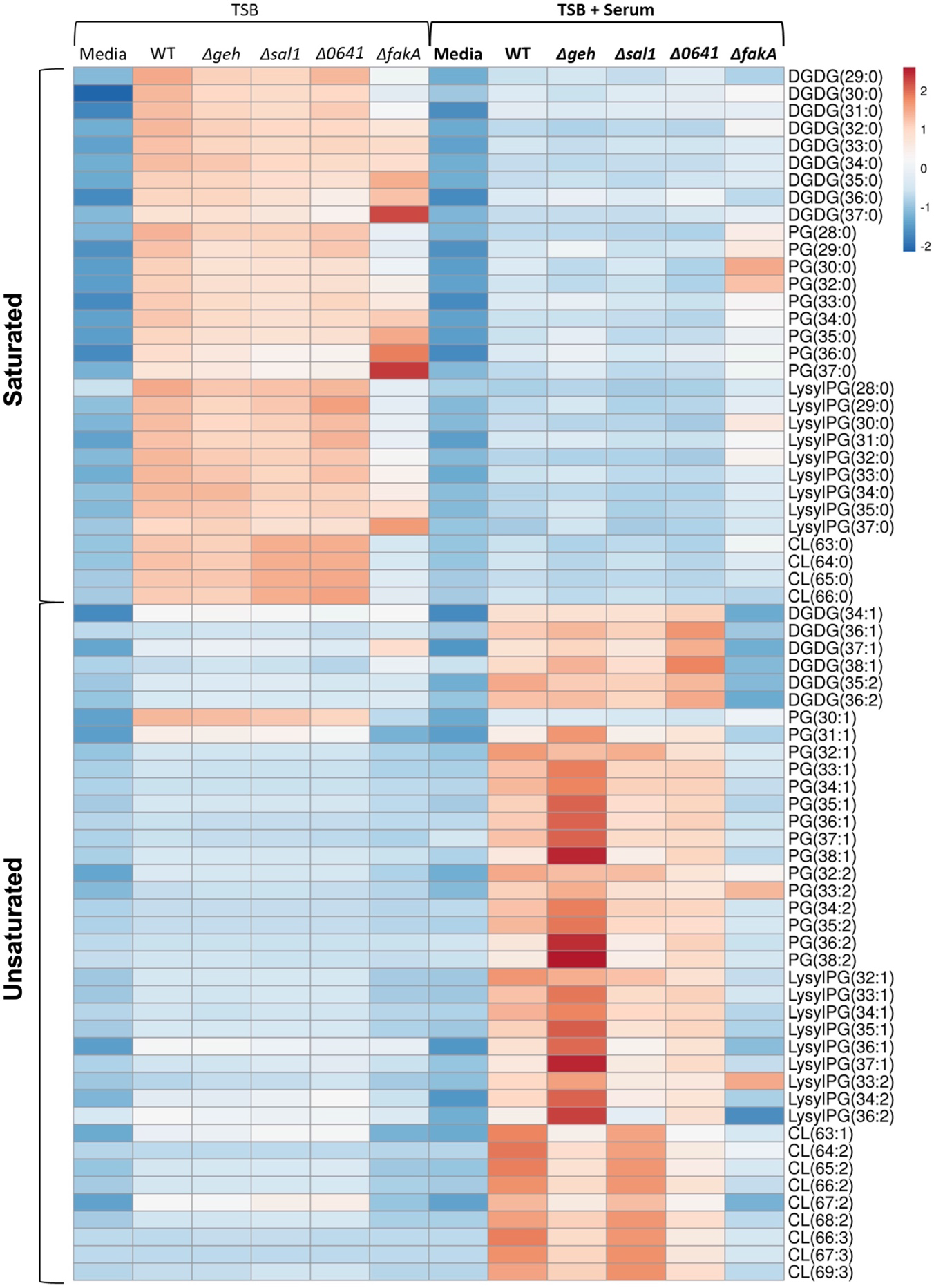
Relative abundances of lipids of WT (USA300 LAC) and *geh*-, *sal1*-, *0641*-, or *fakA*-knockout mutant strains grown in TSB or TSB + 20% human serum. DGDG: diglucosyldiacylglycerol; PG: phosphatidylglycerol; LysylPG: lysyl-phosphatidylglycerol; CL: cardiolipin. Results are row-centered and scaled by unit variance scaling. N = 4 per group.

When WT *S. aureus* was grown in TSB supplemented with 20% human serum, lipid profiles of all membrane lipid classes contained elevated levels of UFAs (such as 33:1, 34:1, 35:1, 36:1, 33:2, 34:2, 35:2, and 36:2) that were absent from strains grown in TSB-only (Figure 2). Linoleic acid (C18:2), palmitic acid (C16:0), and oleic acid (C18:1) comprise the majority of fatty acids found in human serum, along with a slightly lower amount of stearic acid (18:0) and arachidonic acid (C20:4) (42, 43). MS/MS experiments confirmed C18:1 and C18:2 were the dominant UFAs utilized by the WT and lipase mutants (Supplemental Data Set 1). Comparable levels of C20:1 and C20:2 were also observed, suggesting elongation of oleic and linoleic acids by *S. aureus*. When grown in the presence of serum, PG lipids in the WT and lipase KOs with odd-numbered total carbons (e.g, 33 and 35) contain C15:0 as the most abundant acyl side chain while PGs with even-numbered total carbons (e.g., 34 and 36) contain C16:0, instead of C15:0, as a major fatty acid. This pattern was not seen in the Δ*fakA* mutant, however, indicating the increase in C16:0, palmitic acid, likely arose from the serum. As expected, the Δ*fakA* mutant prevented the incorporation of eFAs into the bacterial membrane (Figure 2). This is consistent with previous reports of *S. aureus* incorporating serum-derived UFAs into the bacterial lipids and the necessity of FakA to incorporate eFAs into membrane lipids (3, 7, 8). We noted that Δ*fakA* showed similar intensities to the WT and lipase KOs for PG 32:2, PG 33:2, and LysylPG 33:2 only when grown in TSB-containing human serum. Although individual lipase knockouts did not completely prevent the incorporation of host-derived UFAs, the Δ*geh* mutant exhibited the least UFA abundance overall in DGDGs and CLs. However, Δ*0641* also displayed lower UFA levels than the WT, implying possible overlapping functions exist between the lipases (4, 5). Much higher levels of saturated lipids, especially lipids with saturated chains 30:0 and 32:0, were observed in the Δ*fakA* strain grown in serum, which could indicate upregulation of *de novo* fatty acid synthesis caused by the loss of FakA.

### Substrate specificity of *S. aureus* secreted lipases

To further elucidate the overlapping substrates among the lipases, the WT and Δ*geh*, Δ*sal1*, Δ*0641*, Δ*fakA* mutants were grown in the presence of cholesteryl esters (CEs) and triglycerides (TGs), the major donors of eFAs in serum (3). TSB was supplemented with CE and TG standards containing the unsaturated fatty acids C18:1, C18:2, or C20:4 at a final concentration of 100 μM for each lipid. Comprehensive lipidomics was conducted in the same way as described above (Figures 3 and S1).

**Figure 3.**
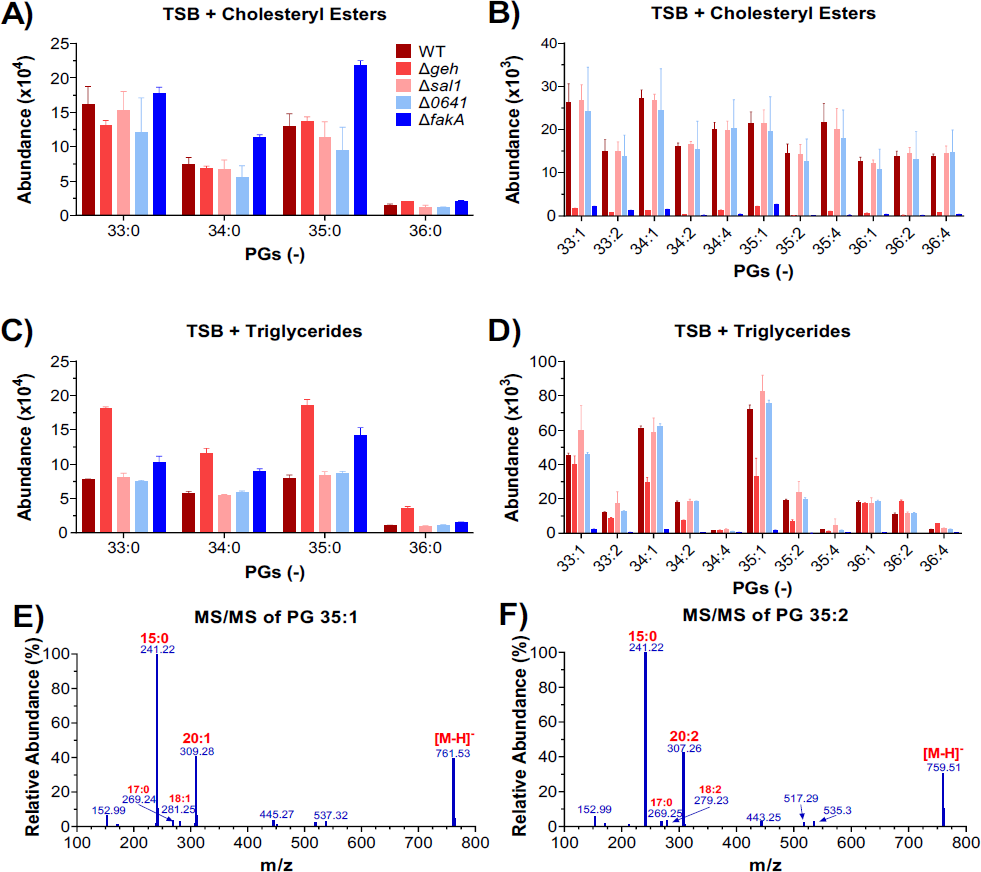
Relative abundance of lipids of WT (USA300 LAC) and *geh*-, *sal1*-, *0641*-, or *fakA*-knockout mutant strains grown in the presence of cholesteryl esters or triglycerides containing C18:1, C18:2, or C20:4 at 100 μM for each lipid. (A) and (B): saturated and unsaturated lipids in the strains grown in the presence of cholesteryl esters, respectively. (C) and (D): saturated and unsaturated lipids in the strains grown in the presence of triglycerides. (E) and (F): MS/MS fragmentation spectra of two unsaturated PGs. N = 4.

When grown in the presence of CEs, the WT, Δ*sal1*, and Δ*0641* strains displayed similar eFA incorporation in PG (Figure 3B), DGDG, LysylPG, and CL (Figure S1) lipid species. Neither the Δ*fakA* nor Δ*geh* strain contained UFAs in any major lipid classes. This suggests that Geh is the lipase responsible for hydrolyzing cholesteryl esters. Elongation of the supplemented CE unsaturated fatty acids was observed in the wild type, Δ*sal1*, and Δ*0641* mutants as evidenced by the presence of C20:1, C20:2, and C22:4 (Supplemental Data in Excel). MS/MS of DGDG and PG lipid species confirmed the fatty acyl composition of 34:1 and 34:2 to be C14:0 and C20:1 or C20:2, 35:1 and 35:2 contained C15:0 and C20:1 or 20:2, and PG 36:4 contained C14:0 and C22:4.

In the presence of TGs, the Δ*geh* strain again had the most significant impact on the incorporation of eFAs (Figure 3C and 3D). However, Δ*geh* did not completely abolish eFA incorporation within PG and LysylPG lipids. Differences in the fatty acid composition of PG 36:1 and PG 36:2 between the wild-type, Δ*sal1*, Δ*0641* strains and Δ*geh* were identified with C18:1 being the most abundant acyl side chain in the Δ*geh* strain and C20:1 for Δ*sal1* and Δ*0641* strains (see Supplemental Data in Excel). Interestingly, increased levels of saturated lipids were observed in Δ*geh*, indicating an upregulation of *de novo* fatty acid synthesis in this lipase KO. Overall, this suggests Geh is the major enzyme hydrolyzing the long-chain triglycerides, but other lipases can also hydrolyze such TGs.

### Human serum albumin as a source of eFAs and its effect on eFA incorporation

Figure 2 showed that eFAs were incorporated into the bacterial membrane when grown in serum, regardless of the lipase knockout, indicating that there may be sufficient amounts of FFAs in the serum, so lipases may not be as necessary in this nutrient-rich environment. FFAs in the bloodstream are typically bound to human serum albumin (HSA), a carrier protein present at high concentrations (35-50 mg/mL) in human blood (44). Albumin concentrations vary throughout the body and sites of infection and decrease with increasing age, highlighting the importance of understanding the effect of HSA on the utilization of serum fatty acids in *S. aureus* (18, 44–46). Here, the WT and Δ*fakA* strains were grown in the presence and absence of fatty acid-containing and fatty acid-free HSA.

We found that fatty acid-containing albumin can indeed serve as a source of eFAs, indicated by the incorporation of UFAs in the WT when grown in the presence of fatty acid-containing HSA (Figure 4A for PGs and Figure S2 for other lipid classes). However, we note that most unsaturated lipids observed when grown in the presence of fatty acid-containing HSA only contain one or two double bonds, much less than those observed when grown in the presence of eFA standards, indicating the majority of fatty acids carried by HSA are mono-unsaturated fatty acids (47, 48). As expected, there were no UFAs incorporated into the membrane lipids with the Δ*fakA* strain or when the WT was grown in the presence of fatty acid-free HSA.

**Figure 4.**
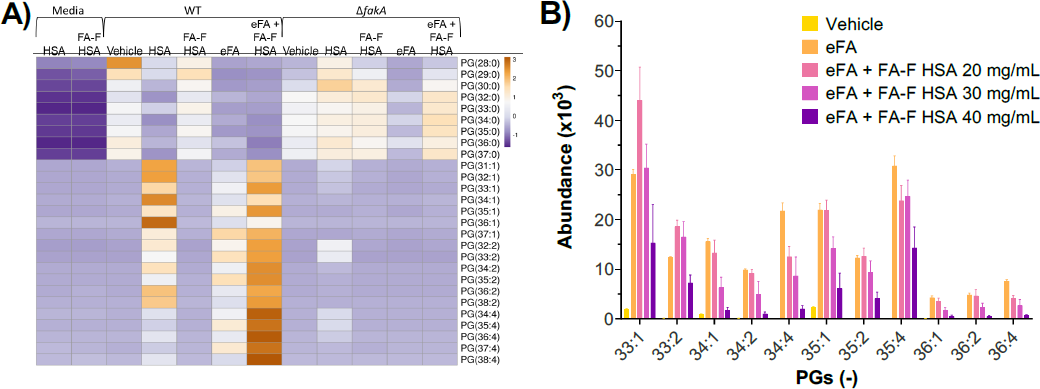
Effect of human serum albumin (HSA) on the incorporation of exogenous fatty acids (eFAs) to WT and *fakA*-knockout strains. (A) Effect of fatty acid-containing HSA and fatty acid-free (FA-F) HSA on the incorporation of eFAs mixture (oleic acid 18:1, linoleic acid 18:2, and arachidonic acid 20:4). (B) The effect of increasing concentrations of FA-F HSA on the incorporation of eFAs. N = 4.

To determine if albumin aids *S. aureus* with incorporating FFAs into the bacterial membrane, the WT and Δ*fakA* mutant were grown in media containing FA-free HSA with the eFA standards: oleic acid (18:1), linoleic acid (18:2), and arachidonic acid (20:4). As seen in Figures 4A and S2, we found that FA-free HSA at 10 mg/mL significantly enhanced the incorporation of UFAs as indicated by the higher levels of unsaturated lipids (Figure 4A). However, concentrations of albumin vary throughout the body, so in a separate experiment, a range of 20 to 40 mg/mL was used. We found that FA-free HSA proportionately decreased the incorporation of UFAs as its concentration increased (Figures 4B and S3). As observed with FA-free HSA at 10 mg/mL, the WT grown with FA-free HSA at both 20 and 30 mg/mL showed greater levels of PG 33:1 and 33:2 than the WT grown with eFAs only. These results suggest that HSA could enhance the utilization of eFAs by *S. aureus* at low concentrations but inhibit the utilization at high concentrations.

### Effect of eFAs on membrane fluidity

Antibiotics, such as daptomycin, have been shown to have increased bactericidal activity against *S. aureus* with incorporated UFAs, which corresponds to increased membrane fluidity, and decreased daptomycin bactericidal activity against *S. aureus* with a high percentage of saturated FAs (25). The membrane fluidity of the WT and Δ*fakA* mutant were assessed at two time points representing the exponential phase (5 hr) and stationary phase (24 hr) of growth with the fluorescent probe 1,6-diphenyl 1,3,5-hexatriene (DHP) in the presence and absence of eFA standards or human serum.

As expected, an increase in membrane fluidity was observed at mid-exponential phase in the WT when grown in the presence of eFAs or serum than without, indicated by a decrease in polarization value (Figure 5A). Comparatively, Δ*fakA* consistently displayed a significantly more rigid membrane compared to the WT in eFAs (*P* < 0.05) and in serum (*P* < 0.05). This is consistent with incorporation of UFAs into the *S. aureus* membrane. However, the Δ*fakA* mutant also displayed overall increases in membrane fluidity in conditions compared to growth in TSB-only during mid-exponential phase (Figure 5A). Although UFAs are not incorporated in the Δ*fakA* mutant, it is possible that the presence of eFAs could lead to a change in endogenous fatty acid synthesis regulation, including synthesis of BCFA, resulting in a more fluid membrane overall. In contrast to the mid-exponential phase of growth, no significant differences in fluidity were found between the WT and Δ*fakA* mutant during the stationary phase (Figure 5B), which may result from the varied lipid composition with growth phases (8, 53). Little difference was displayed between strains in TSB only and TSB containing eFA standards, but both strains were more fluid in the presence of human serum, indicating the impact of the nutritional environment on responses within the membrane composition.

**Figure 5.**
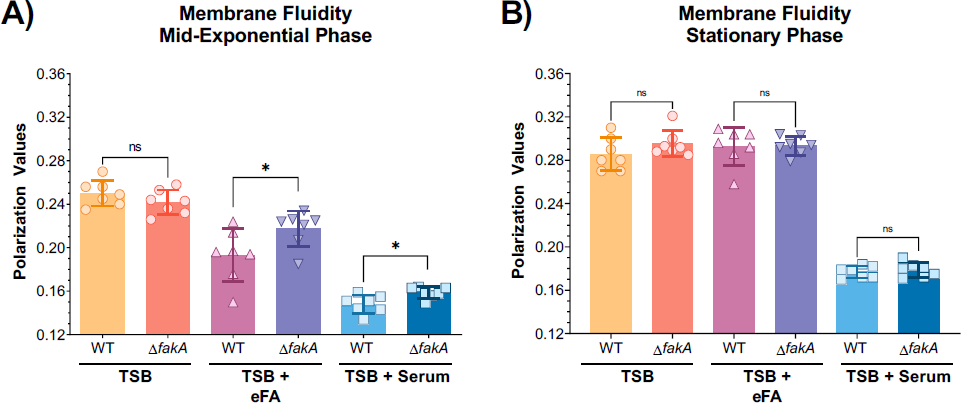
Membrane fluidity of WT and *fakA*-KO (Δ*fakA*) strains grown to the mid-exponential phase (A) or stationary phase (B) in the presence of eFA standards (18:1, 18:2, and 20:4) or 20% human serum. N = 4.

### AFN-1252 enhances UFAs with or without eFA source

An attractive target for drug discovery is the FASII pathway in *S. aureus.* AFN-1252 is an inhibitor that targets the FabI enzyme, an enoyl-acyl carrier protein (ACP) reductase that is essential in the final elongation step of FASII (31, 54, 55). We hypothesized that the FASII inhibitor would enhance the incorporation of eFAs due to suppression of endogenous FA synthesis. Thus, *S. aureus* was grown in the presence of AFN-1252, eFAs, or a combination of both. Exposure of *S. aureus* to AFN-1252 and eFAs resulted in bacterial membrane composed predominantly of UFAs (Figure 6), confirming promotion of eFA incorporation by AFN-1252 (10).

**Figure 6.**
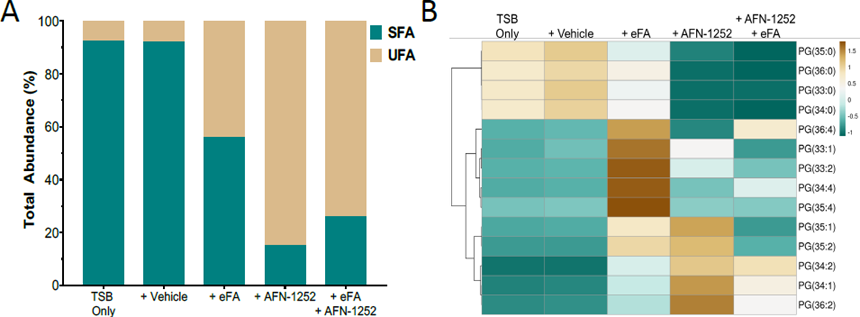
Effect of AFN-1252 on the incorporation of eFA standards containing fatty acids 18:1, 18:2, and 20:4. (A) Comparison of the sum of all saturated and unsaturated lipids; (B) comparison of individual PGs under various conditions. N = 3-4.

Interestingly, the UFA content in the WT grown with only AFN-1252 in the absence of eFAs also increased although displaying a different lipid profile from that of eFA only group (Figure 6 and S4). Upon MS/MS fragmentation, these UFA-containing lipids exhibited different patterns from those grown in the presence of eFAs, mostly containing fatty acids with one double bond. As also seen in prior experiments when *S. aureus* is exposed to eFAs, PG 33:1 was found to be composed of C15:0 (241 *m/z*) and C18:1 (281 *m/z*), but in the presence of AFN-1252 only, PG 33:1 was found to be composed of C14:0 (227 *m/z*) and C19:1 (295 *m/z*) (Figure 7). This is not surprising as AFN-1252 inhibits FabI, which reduces a double bond to a saturated carbon-carbon bond in the FASII cycle, indicating possible accumulation of the ACP intermediate (55). PG 33:2 in *S. aureus* with AFN-1252 contained C14:1, C19:1 (295 *m/z*), C16:1 (253 *m/z*) and C17:1 (267 *m/z*), further suggesting accumulations of the unsaturated ACP intermediate (Figure 7B). Such fatty acid compositional changes reveal a different aspect of the mechanism of action of AFN-1252, which warrants further investigation in the future.

**Figure 7.**
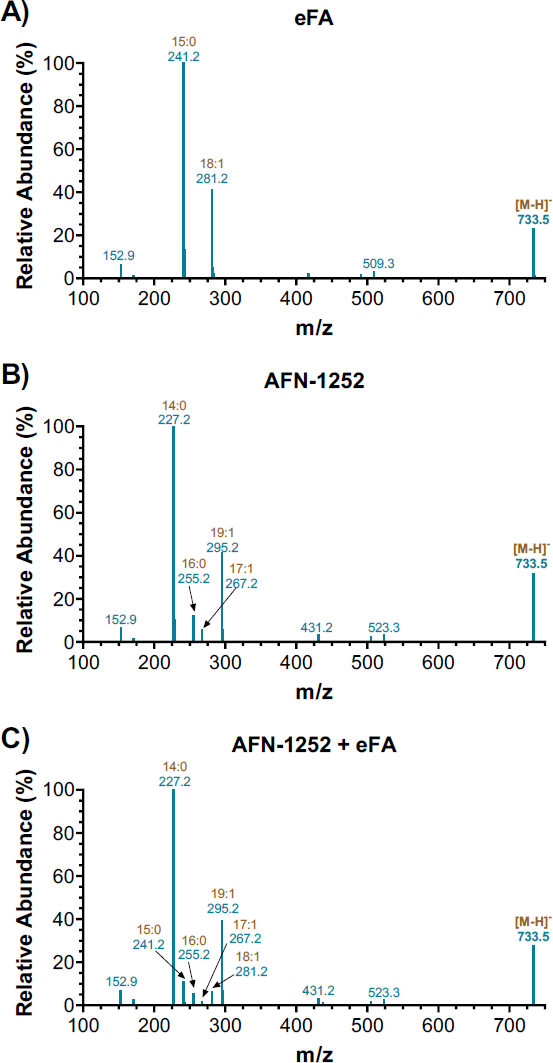
Fatty acid composition of PG 33:1 informed by MS/MS fragmentation of the parent [M+H]^-^ ion from WT grown in the presence of (A) eFA only, (B) AFN-1252 only, or (C) AFN-1252+eFA.

### Effect of eFAs on ROS formation

When *S. aureus* was grown with exogenous fatty acid sources, host-derived fatty acids were incorporated into the membrane, resulting in increased levels of PUFAs (Figures 2, 3, and 4) and growth inhibition by UFAs (Figure 8A). PUFAs such as linoleic acid (18:2), a major UFA found in human skin, and arachidonic acid (20:4), which is released in humans during inflammatory responses, have been shown to be toxic to the bacteria and kill through lipid peroxidation (35, 36). Reactive oxygen species (ROS), produced by phagocytes in PUFA-rich environments, also play an integral role in bacterial killing by oxidative damage (56, 57). To examine the effect of incorporated eFAs on ROS formation in the bacterial cells, the WT and Δ*fakA* mutant were grown with and without eFA standards, and ROS production was measured using the fluorogenic dye, H_2_DCFDA. We observed a significant increase of ROS in the WT strain when the measurements were taken in an eFA-rich environment (Figure 8B). Small increases in ROS formation were also observed in Δ*fakA* mutant, but not as significant as in the WT strain. This suggests that the incorporation of PUFAs into the membrane lipids is necessary to increase oxidative stress and enhance their killing activity.

**Figure 8.**
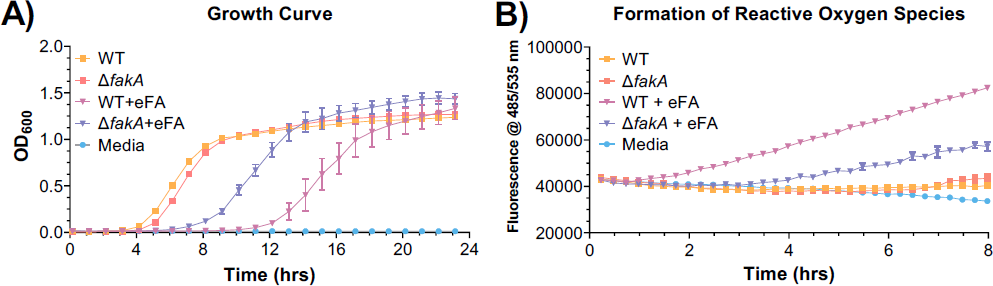
(A) Growth curve and (B) formation of reactive oxygen species in WT and fakA-KO strains in the absence or presence of the eFA standard mixture containing fatty acids 18:1, 18:2, and 20:4.

## DISCUSSION

### Role of lipases in the utilization of serum lipids by *S. aureus*

Although *S. aureus* is known to utilize serum lipids and is thought to depend on Geh to incorporate eFAs from lipoproteins, comprehensive lipidomic studies on the role of bacterial lipases and their substrate specificity on cholesteryl esters have not yet been performed (3, 4, 20, 58). We found that the incorporation of fatty acids from cholesteryl esters required Geh, but not Sal1 and 0641 (Figure 2). On the other hand, none of the lipase mutants grown in the presence of TGs showed a complete lack of UFA incorporation; however, UFAs were decreased in the Δ*geh* mutant compared to Δ*sal1* and Δ*0641.* This is consistent with previous studies that observed a *geh* mutant could still incorporate some UFAs into PG lipids in the presence of human low-density lipoproteins (4). It is likely that Sal1 or 0641 can hydrolyze FAs from TGs to compensate for the absence of Geh. PUFA-containing lipids were not seen at significant levels, whereas monounsaturated lipid species were abundant, implying that the 20:4 PUFA is not preferentially utilized from TGs. Thus, our data suggest that Geh is essential for hydrolyzing UFAs from CEs, whereas other lipases have overlapping functions to release fatty acids from TGs.

### Concentration-dependent effect of human serum albumin on eFA incorporation

We determined that in addition to serum lipoproteins, human serum albumin can serve as a source of eFAs for the bacteria, primarily supplying oleic and linoleic acid (Figure 4A). Although a previous report demonstrated that albumin could sequester exogenous oleic acid from *S. aureus,* preventing the inactivation of the antibiotic daptomycin, that study used fatty acid-free HSA at 10 mg/L (18). Furthermore, we observed that eFA utilization by *S. aureus* had an inverse relationship with albumin concentration, where lower HSA levels promoted FFA incorporation whereas higher levels reduced incorporation. Hypoalbuminemia, diagnosed at albumin levels <35 mg/mL, has recently been significantly associated with increased risk and adverse outcomes of deep musculoskeletal *S. aureus* infections (49, 50). Our findings of albumin concentration affecting eFA incorporation corroborate virulence pathways by which the bacteria utilize host fatty acids to promote survival during infection and tolerate antibiotic treatments (4, 10, 27). Although all lipid species displayed an overall decreasing abundance pattern with increasing albumin concentration, PG 15:0/20:4 levels remained comparatively high at 40 mg/mL, which may be a result of albumin preferentially binding to monounsaturated fatty acids, therefore leaving PUFAs such as arachidonic acid (20:4) and linoleic acid (18:2) more readily available.

### Cell membrane fluidity changes

As expected from incorporating host-derived fatty acids into its phospholipids, the membrane fluidity of *S. aureus* increased in eFA-rich environments (Figure 5). Consistent with previous studies of the Δ*fakA* mutant grown with oleic acid, Δ*fakA* had a significantly more rigid membrane at the mid-exponential phase than the wild-type due to its lack of ability to incorporate eFAs (8, 27). On the other hand, the fluidity of Δ*fakA* strains also increased overall upon eFA and serum treatment (Figure 5A). This provides evidence that differences in membrane fluidity are not entirely due to eFA incorporation, instead suggesting that these environments signal for altered endogenous fatty acid metabolism and composition (8, 61), such as the production of branched-chain fatty acids.

### FASII modification and eFA utilization with AFN-1252 exposure

Therapeutic value of FASII inhibitors remains in debate, as *S. aureus* can bypass suppressed endogenous fatty acid synthesis by utilizing eFAs (9, 11, 18). Lipidomics of *S. aureus* grown with AFN-1252-only revealed a significant increase in the proportion of UFAs with abnormally long chains (C19:1) and phospholipids with various fatty acid combinations (C14:1, C16:1, C17:1, or C19:1), suggesting accumulation of the acyl-ACP intermediate at the inhibited FabI step (9, 55, 64). In the presence of eFAs and AFN-1252, the bacteria indeed incorporated more eFAs than eFAs alone, but the overall UFA content is lower than when treated with AFN-1252 only (Figure 6). This data indicates *S. aureus* preferably continued to initiate new acyl chains, leading to intermediate accumulation, rather than completely favor FASII bypass with eFA; however, preferred pathways and adaptive mechanisms differ based on experimental conditions such as fatty acid sources or FASII inhibitor concentrations (9, 10, 55, 63, 65). AFN-1252 has demonstrated promising synergistic effects when combined with daptomycin by blocking decoy phospholipid release or bacterial growth (18, 65). We speculate that the increased UFA ratio of *S. aureus* in the presence of AFN-1252 could also contribute to enhanced daptomycin activity, as daptomycin targets specific fluid areas of the membrane (15, 25, 66).

To summarize, using comprehensive lipidomics and genetic KOs, this work demonstrated the importance of various *S. aureus* lipases in the utilization of host-derived CEs and TGs, identified a surprising role of HSA as a buffer of eFAs, and revealed an underappreciated biological consequence of the FASII inhibitor AFN-1252, all of which could lead to new approaches to enhance *S. aureus* killing in a host environment.

## MATERIALS AND METHODS

### Bacterial strains and growth conditions

Studies were conducted using the USA300 LAC wild-type (WT) strain of *Staphylococcus aureus*, along with isogenic *Δgeh*, *Δsal1*, *Δ0641*, and Δ*fakA* mutants. *Δgeh*, *Δsal1*, and *Δ0641* mutants were generated as previously described (5). To generate a *ΔfakA* mutant, five hundred-fifty base pair regions of homology upstream and downstream of the *fakA* open reading frame (SAUSA300_1119) were amplified from WT *S. aureus* genomic DNA using primer pairs *fakA*-SOE-1 (CCCGGTACCGGTGATTTAAGCGTAAGTCA) and *fakA*-SOE2 (GGTAGTTTTTTATTTTAAATTTTTCAAGTTGTCCTCCT) or *fakA*-SOE3 (AGGAGGACAACTTGAAAAATTTAAAATAAAAAACTACC) and *fakA*-SOE4 (CCCGAGCTCACCTTTAACAGTTATAGTTTG). The resulting amplicons were used in a splicing by overlap extension (SOE) PCR along with primer pair *fakA*-SOE-1 and *fakA*-SOE4. The final amplicon was subcloned into the pIMAY plasmid after digestion with restriction endonucleases KpnI and SacI (73). Allelic replacement was carried out as previously described (74). This series of knockouts (KOs) target individual lipases or FakA. Each strain was grown to stationary phase in triplicate in 1 mL of tryptic soy broth (TSB) at 37℃ with shaking for 24 hours in Eppendorf tubes. For human serum treatments, TSB was supplemented with 20% heat-treated pooled gender human serum (BiolVT; Hicksville, NY). To determine lipase substrate specificity, the WT and KOs were grown in the presence of pure cholesteryl ester and triglyceride lipid standards found in serum, containing the fatty acid mix: C18:1, C18:2, and C20:4 (Nu-Chek Prep, Inc., Elysian, MN) in ethanol each at 100 μM in TSB. To determine the effect of albumin on eFA sources, the WT and Δ*fakA* mutant were grown with fatty acid-containing and fatty acid-free HSA (Sigma-Aldrich, St. Louis, MO) at 10-40 mg/mL in TSB. To determine the effect of AFN-1252 on eFA incorporation and FASII pathway modifications, the WT was grown in the presence of AFN-1252 (MedChemExpress LLC, Monmouth Junction, NJ) at 0.025 μM in TSB.

### Lipidomics analysis

Cultures were pelleted by centrifugation, washed by resuspension and centrifugation in phosphate-buffered saline (PBS), and dried in a vacuum concentrator. Total lipids were extracted by the method of Bligh and Dyer (65). Dried extracts were reconstituted in 2:1 acetonitrile-methanol. Extracts were analyzed by hydrophilic interaction liquid chromatography (HILIC) coupled with ion mobility-mass spectrometry (IM-MS). Chromatographic separations were carried out with a Phenomenex Kinetex HILIC column (50 x 2.1 mm, 1.7 μm) on a Waters Acquity FTN UPLC (Waters Corp., Milford, MA) (38). The solvent system consists of mobile phases (A) 95% acetonitrile/5% water with 5 mM ammonium acetate and (B) 50% acetonitrile/50% water with 5 mM ammonium acetate. A flow rate of 0.5 mL/min was used with the following linear gradient conditions: 0-0.5 min, 100% A; 2 min, 90% A; 3.5-4 min, 70% A; and 4.5-6 min, 100% A. Injection volumes were 5 μL for both positive and negative modes. CCS calibration was created with phosphatidylcholine and phosphatidylethanolamine CCS standards as previously described (38). IM-MS analysis was performed on a Waters Synapt XS HDMS (Water Corp., Milford, MA) in both positive and negative ionization modes as described previously (wave velocity, 500 m/s; wave height, 40 V) (37, 38). Additional targeted MS/MS experiments were performed with a collision energy ramp of 30-45 eV to determine FA contents of selected DGDG (positive mode) and PG (negative mode) lipid species.

### Data analysis

Data alignment and peak detection were performed in Progenesis QI (Nonlinear Dynamics; Waters Corp., Milford MA) with normalization to all compounds. Retention time calibration and lipid identification were calculated with the Python package, LiPydomics (66). Multivariate statistics were created through LiPydomics and ClustVis (66, 67). MS/MS analysis and identification of the most abundant FAs was performed in Skyline utilizing a targeted lipid library generated with LipidCreator (68, 69).

### Cell membrane fluidity assay

The WT and Δ*fakA* mutant strains were grown to mid-exponential and stationary phase in 5 mL of TSB at 37 ℃ with shaking in Falcon tubes. Each strain was grown in the presence and absence of 20% human serum (v/v) or the fatty acid mix: oleic acid (18:1), linoleic acid (18:2), and arachidonic acid (20:4) (Nu-Chek Prep, Inc., Elysian, MN), each at a final concentration of 100 µM. Cultures were pelleted by centrifugation, washed, and resuspended in normal saline at a McFarland reading of 0.9. Cell membrane fluidity was measured by polarizing spectrofluorometry using a BioTek Synergy H1 plate reader (BioTek Instruments, Winooski, VT) with the fluorescent probe, 1,6-diphenyl-1,3,5-hexatriene (DPH).

### Growth Curves

Overnight cultures of the WT and Δ*fakA* strains were diluted 1:100 in TSB for growth curve measurements. Cells were added to a Costar 96-well flat-bottom microplate and grown with FFA standards. Growth was monitored at 600 nm using a BioTek Synergy H1 plate reader (BioTek Instruments, Winooski, VT) set at 37 ℃ with continuous, double orbital shaking.

### Reactive oxygen species measurements

The WT and Δ*fakA* mutant strain were grown to stationary phase in 7 mL of MHB50 at 37 ℃ with shaking for 24 hours in Falcon tubes. Both strains were grown in the presence and absence of the fatty acid mix: oleic acid (18:1), linoleic acid (18:2), and arachidonic acid (20:4) (Nu-Chek Prep, Inc., Elysian, MN), each at a final concentration of 100 µM. Cultures were pelleted by centrifugation, resuspended in 7 mL MHB50 containing the fluorogenic dye, 2’,7’-dichlorodihydrofluorescein diacetate (H_2_DCFDA) at a concentration of 10 µM, and incubated for 45 minutes at 37℃ protected from light. Cultures were pelleted by centrifugation, washed with saline, and resuspended in 7 mL of MHB50. Cells were added in triplicate to a black Nunc 96-well flat-bottom microplate in the presence or absence of the fatty acid mix with a final volume of 200 µL. Reactive oxygen species were measured by fluorescence readings (λ excitation=485 nm, λ emission=535 nm) using a BioTek Synergy H1 plate reader set at 37 ℃ for 8 hours.

## ACKNOWLEDGEMENTS

This work was supported by NIH grant R01AI136979 (to LX and BJW). FA acknowledge the support from NIH Grant R01AI120994 and Burroughs Wellcome Fund Investigators in the Pathogenesis of Infectious Disease Award (1019120.01).

## Notes

### Competing Interest Statement

The authors have declared no competing interest.

### Summary of Updates

Adding supplemental materials.

