## Supplemental Figures for "Elucidating the Impact of Bacterial Lipases, Human Serum Albumin, and FASII Inhibition on the Utilization of Exogenous Fatty Acids by *Staphylococcus aureus*"

Running Title: Exogenous fatty acids utilization by *S. aureus*

Present addresses:

[#]Dylan H. Ross, Pacific Northwest National Laboratory, Richland, Washington, USA

[#]Nathaniel K. Ashford, University of Washington School of Medicine, Seattle, Washington, USA

[#]Francis Alonzo III, Department of Microbiology and Immunology, University of Illinois- College of Medicine at Chicago, Chicago, Illinois, USA

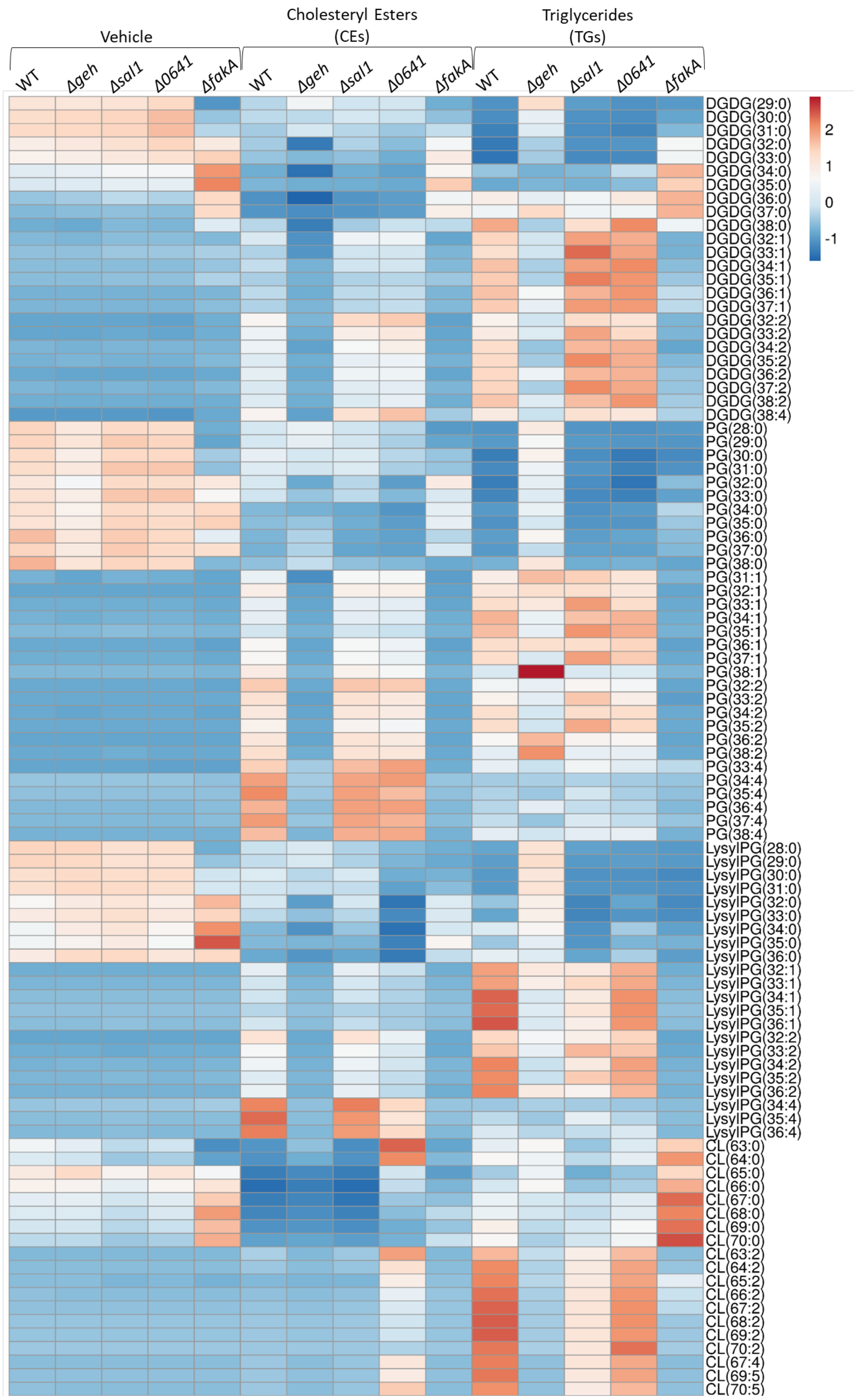


Figure S1. Relative abundances of lipids of WT (USA300 LAC) and *geh*-, *sal1*-, *0641*-, or *fakA*-knockout mutant strains grown in TSB or TSB + cholesteryl esters (CEs) or triglycerides (TGs) containing C18:1, C18:2, or C20:4 at 100 µM for each lipid. Results are row-centered and scaled by unit variance scaling. N = 4 per group.

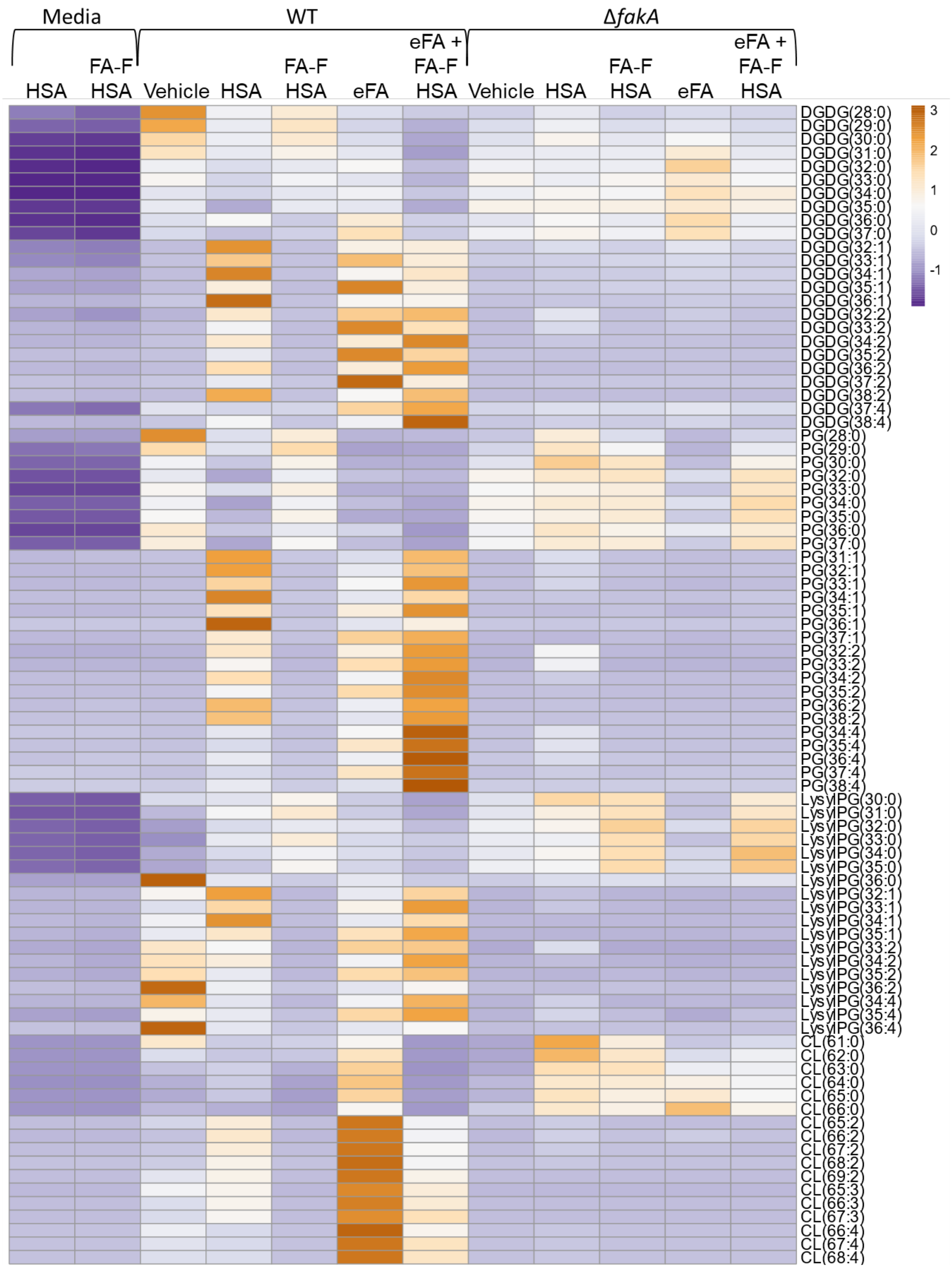


Figure S2. Effect of fatty acid-containing HSA and fatty acid-free (FA-F) HSA at 10 mg/mL on the incorporation of eFAs mixture (oleic acid 18:1, linoleic acid 18:2, and arachidonic acid 20:4) into bacterial lipids. Results are row-centered and scaled by unit variance scaling. N = 4 per group.

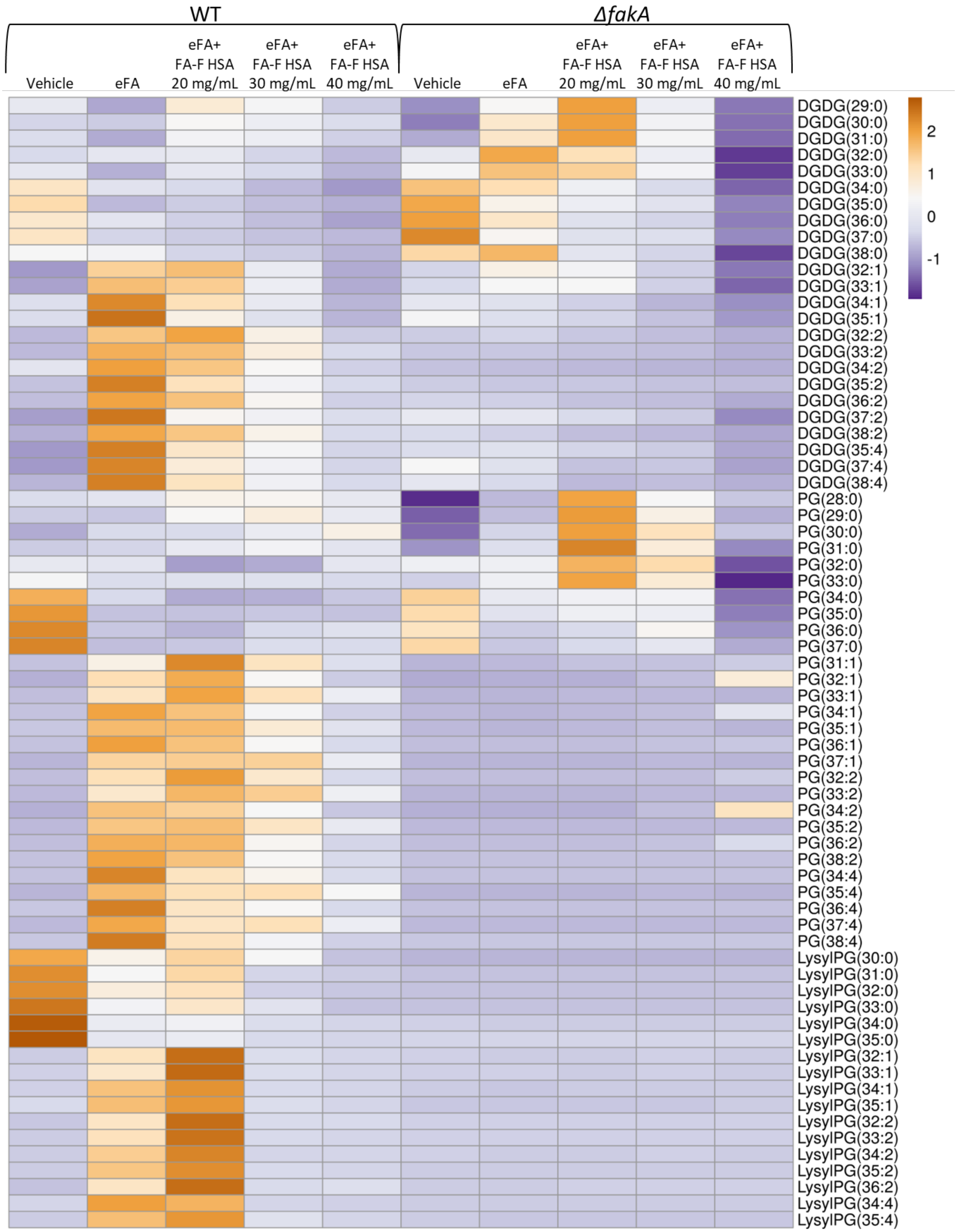

Figure S3. The effect of increasing concentrations of FA-F HSA on the incorporation of eFAs mixture (oleic acid 18:1, linoleic acid 18:2, and arachidonic acid 20:4) into bacterial lipids. Results are row-centered and scaled by unit variance scaling. N = 4 per group.

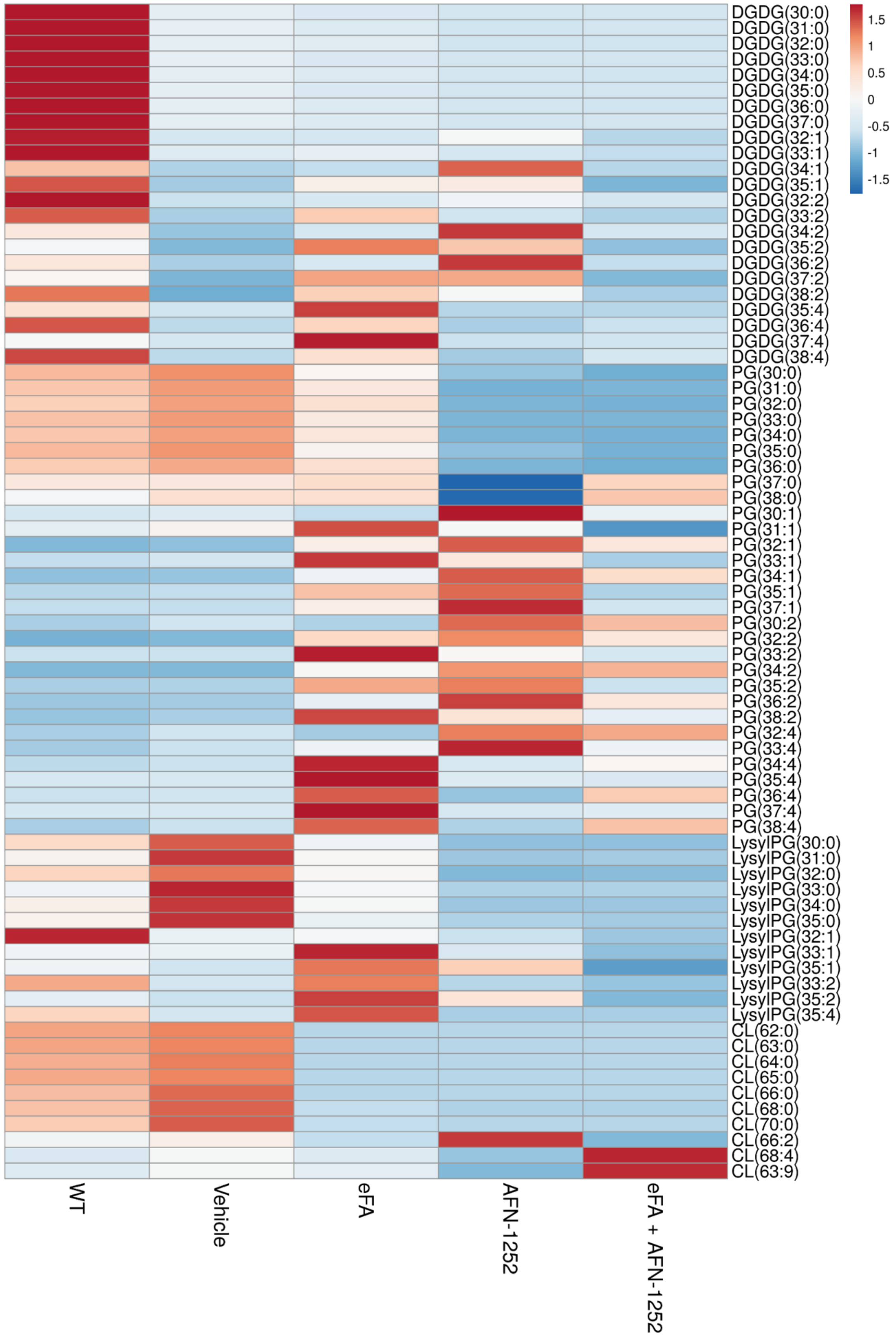


Figure S4. Effect of AFN-1252 on the incorporation of eFA standards containing fatty acids 18:1, 18:2, and 20:4 into various lipid classes in *S. aureus*. Results are row-centered and scaled by unit variance scaling. N = 3-4 per group.
